# Rapid and recent evolution of LTR retrotransposons drives rice genome evolution during the speciation of AA- genome *Oryza* species

**DOI:** 10.1101/086041

**Authors:** Qun-Jie Zhang, Li-zhi Gao

**Affiliations:** Plant Germplasm and Genomics Center, Kunming Institute of Botany, the Chinese Academy of Sciences, Kunming 650204, China; University of the Chinese Academy of Sciences, Beijing 100039, China; Agrobiological Gene Research Center, Guangdong Academy of Agricultural Sciences, Guangzhou 510640, China

**Keywords:** LTR retrotransposons, *Oryza*, AA- genomes, rice speciation, comparative genomics

## Abstract

The dynamics of LTR retrotransposons and their contribution to genome evolution during plant speciation have remained largely unanswered. Here, we perform a genome-wide comparison of all eight *Oryza* AA- genome species, and identify 3,911 intact LTR retrotransposons classified into 790 families. The top 44 most abundant LTR retrotransposon families show patterns of rapid and distinct diversification since the species split over the last ~4.8 Myr. Phylogenetic and read depth analyses of 11 representative retrotransposon families further provide a comprehensive evolutionary landscape of these changes. Compared with Ty1-*copia*, independent bursts of Ty3-*gypsy* retrotransposon expansions have occurred with the three largest showing signatures of lineage-specific evolution. The estimated insertion times of 2,213 complete retrotransposons from the top 23 most abundant families reveal divergent life-histories marked by speedy accumulation, decline and extinction that differed radically between species. We hypothesize that this rapid evolution of LTR retrotransposons not only divergently shaped the architecture of rice genomes but also contributed to the process of speciation and diversification of rice.

## Introduction

Long terminal repeat (LTR) retrotransposons are major components of plant genome modification and reorganization (Bennetzen, 2000; Jiang and Ramachandran, 2013; Wessler *et al.*, 1995). As one of the longest classes of transposable elements, their abundance makes them an important driver of plant genome size variation (Piegu *et al.*, 2006; Vitte and Panaud, 2005). For instance, the *Arabidopsis thaliana* genome (~157 Mb) has very limited number of LTR retrotransposons, 5.60% (Pereira, 2004), the rice genome (~389 Mb) is comprised of ~22% LTR retrotransposon sequences (Ma *et al.*, 2004), and 74.6% of the maize genome (2,045 Mb) is occupied by LTR retrotransposon elements (Baucom *et al.*, 2009a). Moreover, both LTR reverse transcriptase activity and the host genome together help restrain mechanisms such as the deletion, unequal recombination, and methylation, affecting the overall abundance of LTR retrotransposons (Bennetzen, 2002; Petrov *et al.*, 2000; SanMiguel *et al.*, 1998; Vitte and Panaud, 2003). Differential retrotransposition activity and DNA loss rates affect the half-life of retrotransposon events in different plant species; wheat and barley, for example, were found to have far longer periods of retrotransposon activity when compared to rice (Wicker and Keller, 2007). The nature and dynamic changes of LTR retrotransposons during the speciation process are poorly understood.

The structure of LTR retrotransposons is similar to retroviruses (Xiong and Eickbush, 1990), encoding for two proteins: *gag* and pol. Previously, the position of the reverse transcriptase (RT) gene in relation to the integrase (IN) gene of *pol* was used to classify the retrotransposon families into Ty1-*copia* (PR-IN-RT) and Ty3-*gypsy* (PR-RT-IN), respectively (Coffin, 1997; Eickbush and Jamburuthugoda, 2008). Extensive investigations in diverse plant genomes have shown that at least six ancient Ty1-*copia* and five Ty3-*gypsy* lineages existed before the divergence of monocots and dicots (Du *et al.*, 2010; Wang and Liu, 2008). Recent studies have revealed considerable difference in the proportion of Ty1-*copia* and Ty3-*gypsy* elements among many plants, such as maize (Meyers *et al.*, 2001), *Medicago truncatula* (Wang and Liu, 2008), *Populus trichocarpa* (Cossu *et al.*, 2012), *Orobanche* and *Phelipanche* (Piednoel *et al.*, 2013), consistent with their role in determining the genome size variation. In addition, a large proportion of LTR retrotransposons are comprised of nonautonomous elements (Wawrzynski *et al.*, 2008), the replication of which relies completely, or at least in part, on proteins expressed by other elements elsewhere in plant genomes (Vitte and Panaud, 2005). In the rice genome, for example, *Dasheng* and *RIRE2* were previously characterized as a nonautonomous LTR retrotransposon family and its putative autonomous partner, respectively. Both types of retrotransposon elements have similar patterns of chromosomal distribution and target site sequences (TSD), suggesting that they use the same transposition machinery and are likely co-expressed (Jiang *et al.*, 2002). Individual retrotransposon families usually have their own amplification histories, the majority of which exhibit an increased rate of transposition at different periods during the evolutionary process (Baucom *et al.*, 2009b; Vitte *et al.*, 2007; Wicker and Keller, 2007). Specific LTR retrotransposon families, thus, expand at distinct evolutionary periods, because some families are especially prone to be more active than others until mutated (Estep *et al.*, 2013). Comparisons of closely related plant species are important to refine burst rates, molecular evolution and patterns of LTR retrotransposon changes during and after speciation.

The availability of rice reference genome sequences has offered an unparalleled opportunity to understand the evolution of plant retrotransposons, including retrotranspositional dynamics, the rates of amplification and removal of the LTR retrotransposons as well as natural selection within LTR retrotransposon families in the rice genome (Baucom *et al.*, 2009b; Ma *et al.*, 2004; Vitte and Panaud, 2003). Comparative genomic analyses among multiple divergent plant lineages have added considerable insight into the conservation and evolutionary dynamics of ancient retrotransposon lineages (Jiang and Ramachandran, 2013; Roulin *et al.*, 2009; Wicker and Keller, 2007). To our knowledge, however, little is known about genome-wide patterns of the gain and loss of recently amplified LTR retrotransposons and evolutionary birth-and-death processes of different families during plant speciation. With this regard, comprehensive comparisons of very closely related plant species that span the speciation continuum and diverged close to the period of half-life of LTR retrotransposons can significantly improve the inference precision and sensitivity of LTR retrotransposon evolution.

The genus *Oryza* serves as an ideal group fulfilling the requirement to study the recent evolution of LTR retrotransposons. They comprise approximately 21 wild and two cultivated species, which can be classified into ten distinct genome types (AA, BB, CC, EE, FF, GG, BBCC, CCDD, HHJJ, and HHKK) (Aggarwal *et al.*, 1997; Ge *et al.*, 1999). Among them, *O. australiensis* (EE genome, ~965 Mb) has the largest genome size, nearly doubling its genome size by accumulating over 90,000 retrotransposons (Piegu *et al.*, 2006). On the contrary, *O. brachyantha* (FF genome, ~261 Mb) has the smallest genome size with a limited number of retrotransposons (Chen *et al.*, 2013; Uozu *et al.*, 1997). The AA- genome *Oryza* species, also called as the *Oryza sativa* complex, consist of two cultivated rice species, Asian cultivated rice (*O. sativa)* and African cultivated rice (*O. glaberrima)*, and six wild rice (*O. rufipogon, O. nivara, O. barthii, O. glumaepatula, O. longistaminata, O. meridionalis*), which are disjunctively distributed in pantropical regions of the four continents of Asia, Africa, South America and Australia (Vaughan, 1989; Vaughan *et al.*, 2003). The recent phylogenomic analysis of these eight diploid AA-genome species supports a series of closely spaced speciation events in this genus (Zhu *et al.*, 2014). Previous studies have identified numerous LTR retrotransopson families that were found to have undergone bursts of amplification within the last five million years (Myr) in the *O. sativa* genome (Matyunina *et al.*, 2008). Such a time scale seems older than the earliest divergence time estimated to split from a common AA-genome ancestor ~4.8 Myr (Zhu and Ge, 2005; Zhu *et al.*, 2014).

Here, we perform a genome-wide comparison in a phylogenetic context, and characterize the evolutionary dynamics of LTR retrotransposons across eight completed or nearly finished AA- genomes of the *Oryza* (Zhang *et al.*, 2014). Our study has, for the first time, fully reconstructed the evolutionary history of LTR retrotransposon families in closely related rice species. These data provide a starting point for exploring how evolutionary dynamics of LTR retrotransposons can strongly influence plant genome size variation and genome evolution during the process of recent plant speciation.

## Methods

### Eight genome sequences of *Oryza* AA- genome species

The genomic sequences of *O. sativa* ssp. *japonica.* cv. Nipponbare (Release 7) were downloaded from http://rice.plantbiology.msu.edu. The draft genomes of the other seven AA- genome *Oryza* species of *O. rufipogon, O. nivara, O. glaberrima, O. barthii, O. glumaepatula, O. longistaminata* and *O. meridionalis* were recently sequenced and published (Zhang *et al.*, 2014).

### Annotation and classification of LTR retrotransposon elements

We performed *de novo* searches for LTR retrotransposons against the eight rice genome sequences using LTRSTRUC (McCarthy and McDonald, 2003). False positives caused by long tandem repeats were manually removed by BLAST searches. All intact LTR retrotransposons were classified into Ty1-*copia*, Ty3-*gypsy* and unclassified groups according to the order of ORFs using PFAM (Finn *et al.*, 2008). The RT sequences were retrieved from each retrotransposon element and further checked by homology searches against the published RT genes available from GyDB (http://gydb.org/) (Llorens *et al.*, 2011). They were aligned using ClustalW (Larkin *et al.*, 2007) and manually curated **(Table 1)**. Previous LTR retrotransposon family nomenclature (see **Figure 1** and **Figure 2**) was determined using Blast searches with LTR retrotransposons downloaded from TIGR (Ouyang and Buell, 2004) and Repbase (Jurka, 1998, 2000; Jurka *et al.*, 2005). A homology search of the genome sequence was performed using RepeatMasker (Smit *et al.*). All intact LTR retrotransposon sequences generated by LTRSTRUC (McCarthy and McDonald, 2003) were classified into families using BLASTClust (http://www.ncbi.nlm.nih.gov/Web/Newsltr/Spring04/blastlab.html) and all-to-all BLAST of 5’ LTR sequences, followed by manual inspection (Llorens *et al.*, 2011). The family classification standard was considered acceptable if more than 50% of the 5’ LTRs and sequence identity was greater than 80%. Detailed information of LTR retrotransposon families is provided in **Table S1**.

**Table 1.**
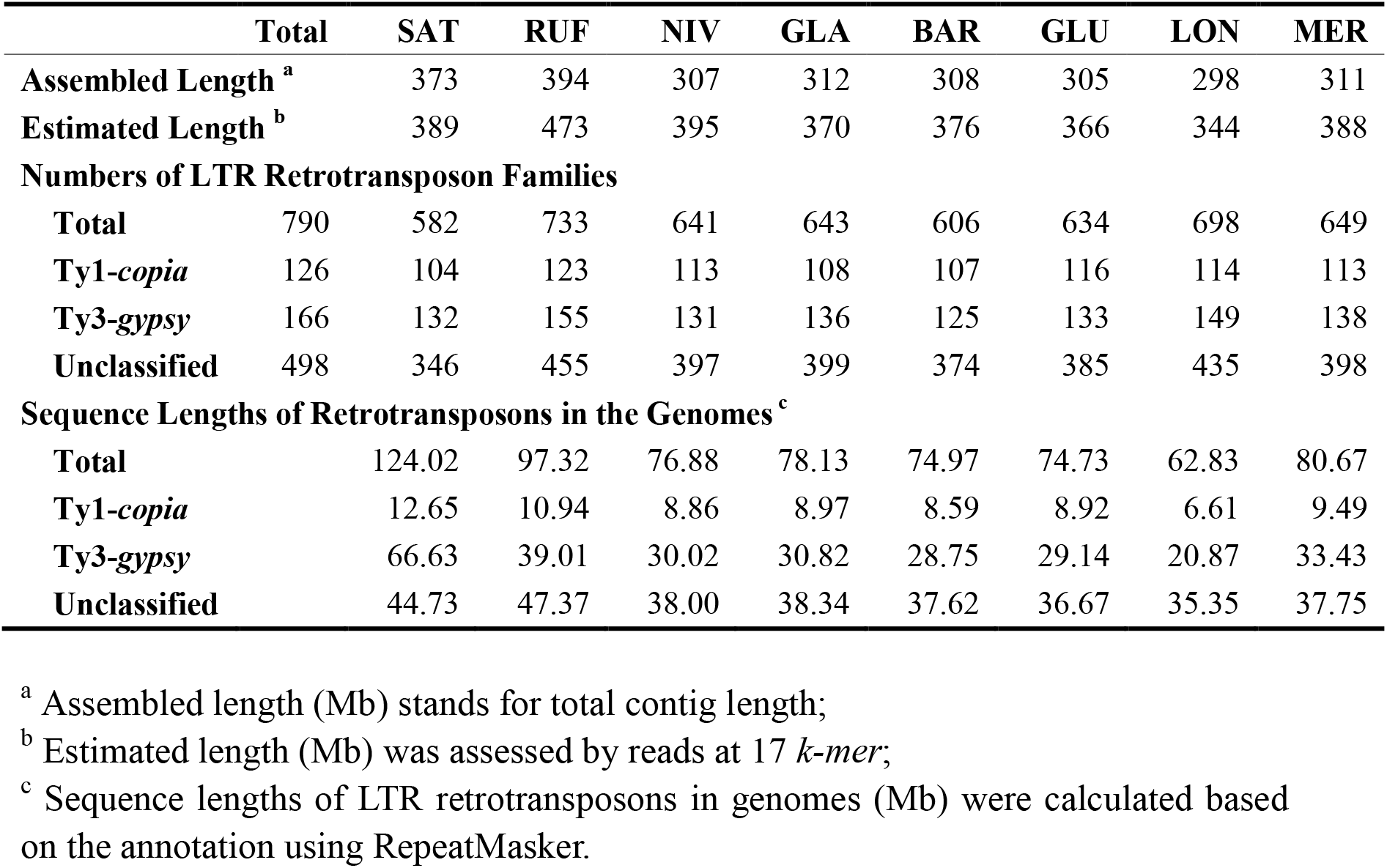
Statistics of the LTR retrotransposons in the eight AA- genome *Oryza* species.

**Figure 1.**
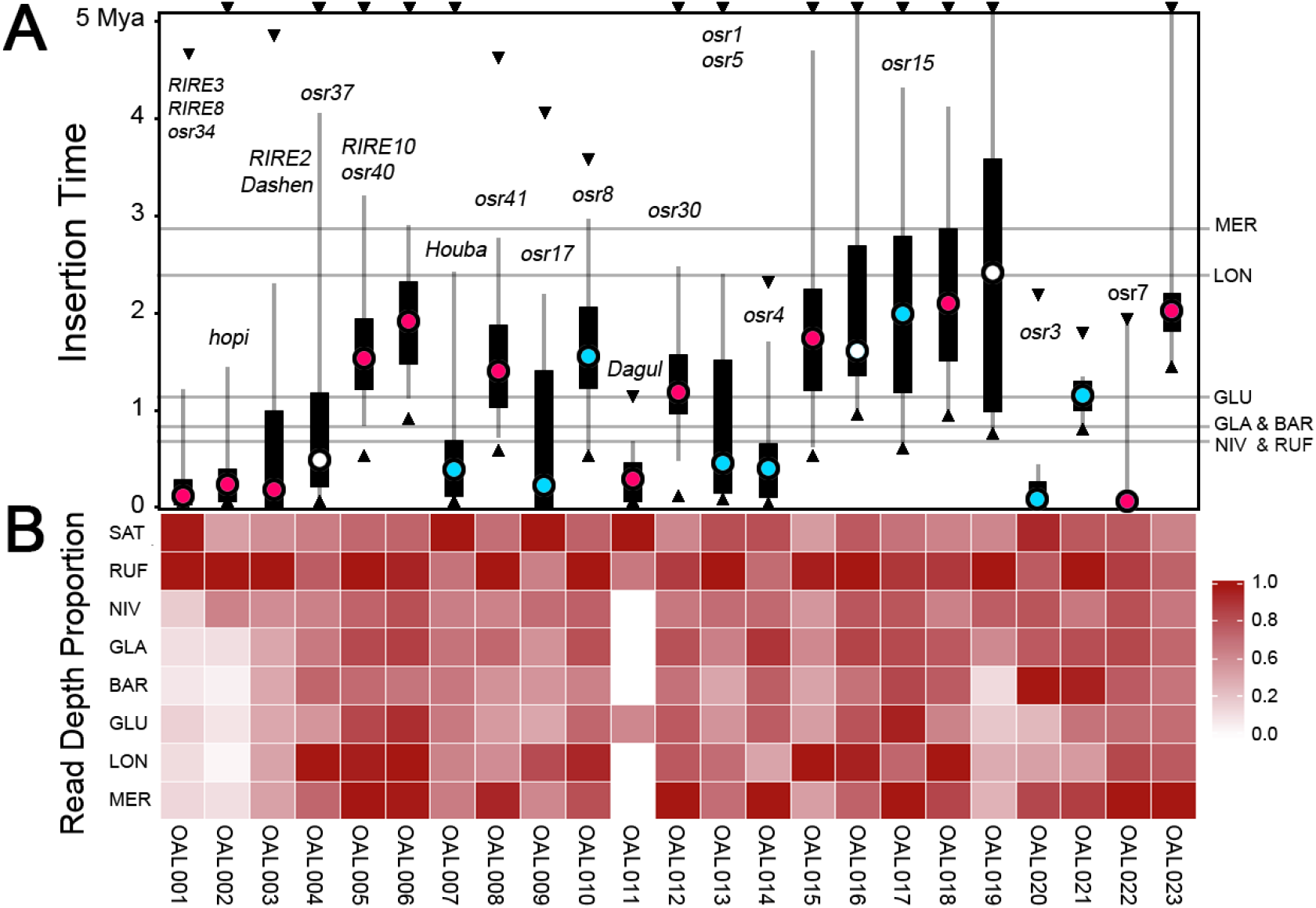
Insertion times and read depth analysis of the top twenty-three LTR retrotransposon families. **(A)** Insertion times of the exemplar LTR retrotransposons from SAT. Black circles indicate mean values, black bars signify 25% to 75% of values, dark grey lines represent 5% to 95% of values, and green circles denote extreme values. The light grey horizontal lines show divergence times between SAT and the other seven species. Those inserted earlier than 5 Myr are set at ~5 Myr. Ty3-*gypsy* (red dots) Ty1-*copia* (blue) elements are distinguished. **(B)** Heatmap of the proportions of LTR read depth compared within each family across the eight rice genomes.

**Figure 2.**
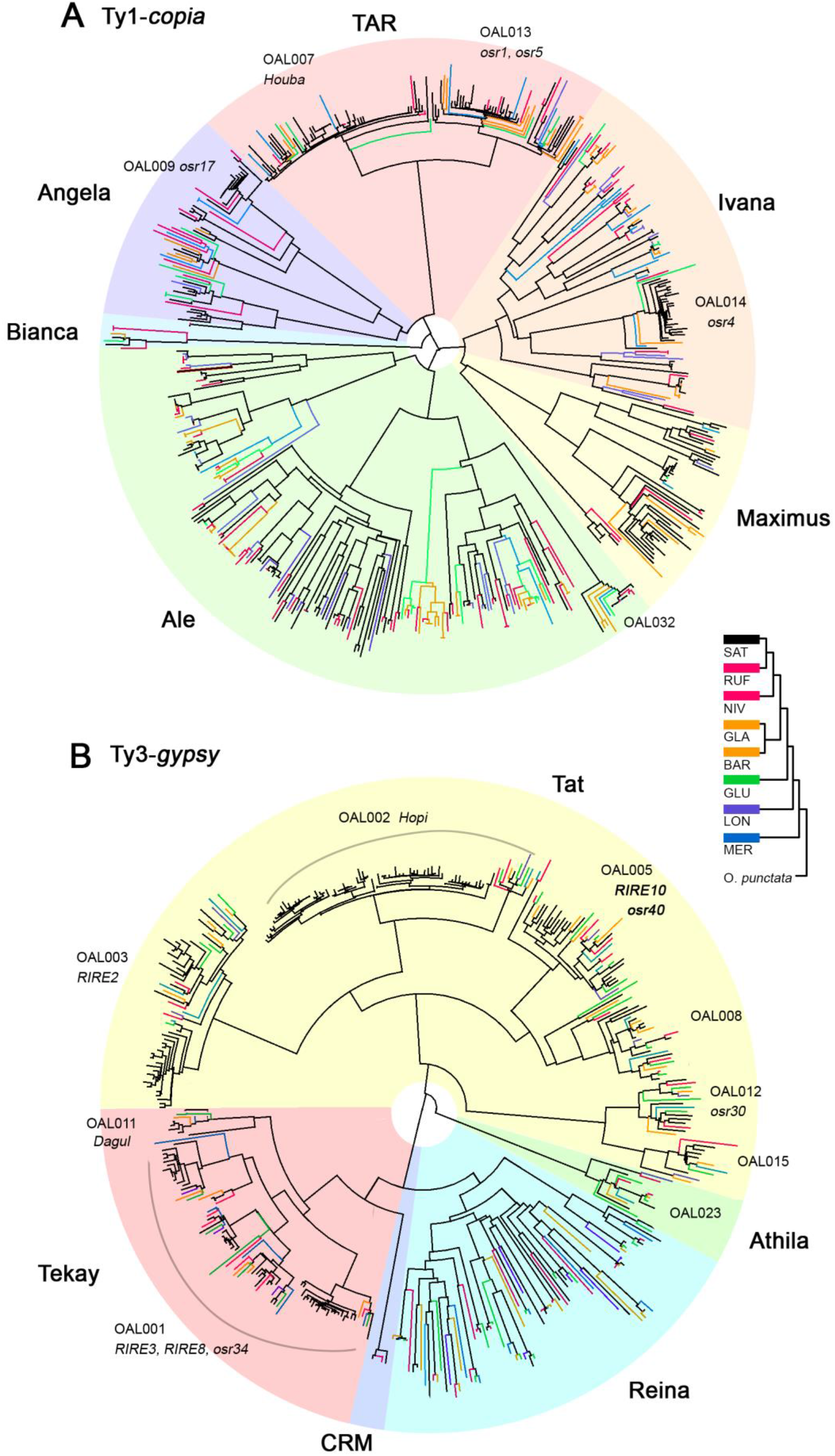
Phylogenetic trees of representative LTR retrotransposon lineages across the eight AA- genome *Oryza* species. Neighbor-Joining and unrooted trees were constructed based on sequences of RT genes for Ty1-*copia* **(A)** and Ty3-*gypsy* **(B).** The backbone of the RT trees and retroelements from SAT are shown in black, while colored branches indicate those from other seven species. LTR retrotransposons from NIV and RUF (the closest to SAT), GLA and BAR (similar divergence time to SAT), GLU, LON and MER are colored in red, orange, green, light blue and dark blue, respectively.

### Dating LTR retrotransposon elements

Dating LTR retrotransposons assumes that the two long terminal repeats (LTRs) are identical when they inserted into the host genome (SanMiguel *et al.*, 1998). The insertion times of intact LTR retrotransposon elements were calculated based on the previously published approach (SanMiguel *et al.*, 1998). The two LTRs of each intact LTR retrotransposon that contains TSD were aligned using ClustalW (Larkin *et al.*, 2007) and their nucleotide divergence was estimated using the baseml module implemented in PAML (Yang, 2007). The insertion times were then computed using: *T* = K/2r. Where: *T*=insertion time; r=synonymous mutations/site/Myr; K=the divergence between the two LTRs. A substitution rate of 1.3×10^−8^ per site per year was used to calculate insertion times (Baucom *et al.*, 2009b; Vitte *et al.*, 2007).

### Phylogenetic analysis

Nucleotide sequences of reverse transcriptase domains (RT) were retrieved from the intact LTR retrotransposon elements of SAT. For the other seven species, RT sequences were also included that were annotated by RepeatMasker (Smit *et al.*), following the guidelines set forth by Xiong and Eickbush (Xiong and Eickbush, 1990). Sequence alignments of amino acid sequences of the RT regions were performed by using ClustalW2 (Larkin *et al.*, 2007) and were adjusted manually. The neighbor joining (NJ) method was used to generate unrooted trees using uncorrected pairwise distances from the sequence alignments with the program MEGA 6 (Tamura *et al.*, 2007). In total, 2,420 Ty3-*gypsy* and 983 Ty1-*copia* RT sequences were extracted to construct phylogenetic trees. For convenience, we only display tree topologies using 414 Ty3-*gypsy* and 447 Ty1-*copia* retroelements (**Figure 2**) after removing highly similar sequences. These Ty3-*gypsy* and Ty1-*copia* RT sequences were classified into 11 lineages, consistent with previous results (Hřibová *et al.*, 2010; Llorens *et al.*, 2009; Vitte *et al.*, 2007; Wicker and Keller, 2007). The RT sequences of SAT were all derived from intact LTR retrotransposons. In the other seven species, besides intact LTR elements, we included partial RT sequences annotated by RepeatMasker (Smit *et al.*).

### Read-depth analysis

To investigate the abundance and evolutionary dynamics of the LTR retrotransposon families we performed read-depth analysis to estimate LTR retrotransposon copy number. The libraries of reference LTRs were constructed using the output from both LTRSTRUC (McCarthy and McDonald, 2003)and RepeatMasker (Smit *et al.*) for each species after removing sequence redundancy using cd-hit-est (Fu *et al.*, 2012; Li and Godzik, 2006) at an identity cutoff of 0.95 (Zhang *et al.*, 2006). Some highly similar genomic regions failed to be assembled for the seven draft AA- genome sequences using Illumina sequencing technology. Approximately 5-fold sequence coverage of Illumina 100 PE reads from each species were randomly sampled and mapped to each reference LTR by SOAPaligner/soap2 (Li *et al.*, 2009). Actual read depths for each LTR were estimated by dividing the depths obtained by the average read-depth for the whole genome. For uncertain reads mapping between LTRs and inner regions, only LTR depths were computed to estimate the abundance of the representative families. Considering that truncated reference LTRs may influence the estimation of mapping depths, LTRs shorter than 150 bp were excluded. We determined that LTRs shorter than 150 bp, in part, belonged to *Ale* of Ty1-*copia*, while others were members of unclassified families; most of these were single-copy families and had been silenced within the last 3 Myr.

## Results and Discussion

### Genome-wide assessment of LTR retrotransposon abundance

To discover the abundance of LTR retrotransposons across all eight AA- genome *Oryza* species, we characterized these elements by using an integrated approach that considers both structure and homology, as described in **Methods**. Besides *O. sativa* ssp. *japonica* cv. Nipponbare genome (abbreviated as SAT) (Release 7, IRGSP, http://www.ncbi.nlm.nih.gov), seven recently completed draft AA- genomes (Zhang *et al*., 2014) were used in this study, including *O. rufipogon* (RUF), *O. nivara* (NIV), *O. glaberrima* (GLA), *O. barthii* (BAR), *O. glumaepatula* (GLU), *O. longistaminata* (LON), and *O. meridionalis* (MER). The order of these species used in this study reflects the topology of phylogenetic tree we recently reconstructed (Zhu *et al.*, 2014). Our retroelement discovery process yielded a total of 3,911 intact LTR retrotransposons in the eight rice genomes after removing ~30 redundant elements. We define an intact retrotransposon element as a copy that has both complete LTR ends, but does not make any statement of whether it encompasses internal insertions or deletions. These intact elements were subsequently clustered into different families using BLASTClust and all-to-all BLAST (Llorens *et al.*, 2011) (**Table S1**). We define a “family” based on 5’ LTR sequence identity. Because LTRs do not encode proteins, they are among the most rapidly evolved sequence regions of the retrotransposons. We consider two retroelements as belonging to the same family if their LTR sequence identity exceeds 80% and they show 50% reciprocal overlap in their lengths. Note that these criteria are somewhat stricter than those reported in other studies (Baucom *et al.*, 2009b; Seberg and Petersen, 2009), as we aimed to detect the variation and divergence of LTR retrotransposon families among these closely related species. As a result, we could classify intact rice LTR retrotransposon elements into a total of 790 families, of which there were 99 multi-member families with more than two intact copies and 160 single-member families in SAT. The remaining 531 families including both single- and multi- members were identified among the other seven non-SAT genomes. This suggests the generation and expansion of a large number of retrotransposon families after the divergence of SAT and non-SAT genomes. Using PFAM (Finn *et al.*, 2008) and tBlastN (1e-10, coverage >=30%) we further grouped them into 126 Ty1-*copia* families comprising 775 intact elements and 166 Ty3-*gypsy* families with 1,803 intact elements. The other 498 families including 1,333 intact elements that lack the *pol* gene were categorized as unclassified families (**Table 1**; **Table S1**). Even though there are fewer Ty3-*gypsy* families than Ty1-*copia*, Ty3-*gypsy* occur more prevalently than Ty1-*copia* elements in these eight rice genomes, as observed on SAT alone and the FF- genome species, *O. brachyantha* (Chen *et al.*, 2013). We operationally named family IDs by the number of the intact elements in SAT combined with the initials of the *Oryza* **A**A- genome of **L**TR retrotransposons (OAL) (i.e. OAL001, OAL002, OAL003, and so on). Of these identified multi-copy families, a total of 31 were previously described, and thus their corresponding family names used in earlier references are also provided in **Table S2**. Our results show that the majority of families, if not all, are shared by all eight rice genomes, but their copy number varied dramatically among the species. More LTR retrotransposon families were identified among non-SAT genomes, despite the higher assembly quality of the SAT reference genome. The data ensures broad LTR retrotransposon representation necessary to study their diversity and evolution. The largest number of LTR retrotransposon families was detected in RUF, followed by LON consistent with known differences in genome size. To investigate the proportion of LTR retrotransposon sequence within these eight AA- genomes, we also annotated their sequence length using RepeatMasker (Smit *et al.*). Although fewer LTR retrotransposon families are detected in SAT, the overall content of LTR retrotransposons in SAT was greater than any of the other seven genomes likely due to the better assemble quality as a reference genome and the inherent difficulty of assembling full length LTR retrotransposons in non-SAT genomes with NGS technology.

To assess retrotransposon expansion and contraction across the eight AA-genome *Oryza* species, we performed a read-depth analysis of all identified retrotransposon families against their own assembled genomes. To test the reliability of read depth to estimate retrotransposon copy number variation, we compared the observed genome copy number of intact elements and LTR sequence read depth in SAT (**Figure S1A**). The results reveal that the number of intact elements significantly correlates with LTR read depth (r = 0.496, *P* < 0.01), and that LTR read depth may serve as a good proxy to evaluate the LTR retrotransposon abundance. Read depth analysis of the most abundant 44 families measured by copy number further showed a significant correlation of SAT LTR read depth and each of the other seven species (**Figure S1B**). Taken together, our results suggest that LTR retrotransposon families experienced rapid diversification after recent spilt of these eight AA- genome *Oryza* species over the past 4.8 Myr.

### Early integration of most LTR retrotransposon families before the split of rice species

When a retrotransposon element integrates into the host genome, the two LTR sequences are assumed to be identical. Thus, we may estimate the insertion times of LTR retrotransposons based on the sequence divergences between LTR pairs. Because the LTR sequences evolve more rapidly than genes, we employed an average substitution rate (r) of 1.3 × 10^−8^ substitutions per synonymous site per year to estimate insertions times (Ma *et al.*, 2004). LTR sequences of the 3,911 complete LTR retrotransposons from the most abundant 44 families were sampled to calculate their integration times (**Table S1**). To trace when these retrotransposon elements came into the eight AA- genomes, we searched and annotated overall features of the top 23 of these 44 retrotransposon families. In total, 2,213 complete retrotransposon elements were dated by LTR identity and projected onto a phylogenetic tree of the eight AA-genome *Oryza* species (**Figure 1**). Our results show that almost all high-copy families, except for OAL011, could be detected and the earliest insertion events for 18 families occurred before the AA- genome *Oryza* species diverged. The other five retrotransposon families appear younger, but may have lost more ancient LTR retrotransposon signatures due to a high turnover or interlocus gene conversion that destroy or homogenize LTR retrotransposon structure. Others such as OAL011 likely represent recently expanded retrotransposon families.

Phylogenetic trees of 11 representative retrotransposon lineages were constructed based on conserved reverse transcriptase (RT) domains for both Ty1-*copia* and Ty3-*gypsy* elements (**Figure 2**). Our results showed that, besides the majority of newly identified families in this study, the previously characterized LTR retrotransposon families including Ty1-*copia* and Ty3-*gypsy* could be found in all eight AA- genome *Oryza* species (**Figure 2**) (Du *et al.*, 2010; Wang and Liu, 2008; Wicker and Keller, 2007), suggesting their early integration into the common ancestral genome. Of these 790 families, we identified 374 solo and full-length LTRs that were shared amongst these rice species. Only 26 that belong to single-copy families may be species-specific, while others were the members of unclassified retrotransposon families (**Table S1**). The majority of LTR retrotransposon families recently came to the most recent ancestral genome before the divergence of all eight AA- *Oryza* genome.

### Evolutionary landscape of Ty1-*copia* and Ty3-*gypsy* retrotransposons

Phylogenetic analyses of 11 representative retrotransposon lineages further show the evolutionary dynamics of rice LTR retrotransposons, including Ty1-*copia* (*TAR, Ivana, Maximum, Ale, Bianca* and *Angela*) (**Figure 2A**) and Ty3-*gypsy* (*Tat, Athila, Reina, CRM* and *Tekay*) (**Figure 2B**). Sequence lengths were calculated for each retrotransposon lineage using Repeatmasker (Smit *et al.*) and then compared across these rice genomes to characterize their content and contribution to genome size variation (**Figure S2A and B**). Since the genome assembly quality may affect genome annotation, we specifically generated the histograms of total length in SAT as a control (**Figure 3A and B**), revealing a consistent pattern in comparison with the other seven non-SAT draft genomes. Considering the difficulty of assembling newly amplified retrotransposons in these non-SAT genomes due to high sequence similarity, we complemented this analysis by LTR read depth estimates (**Figure 3C and D**). The integrated data provide a more comprehensive framework for assessing how Ty1-*copia* and Ty3-*gypsy* retrotransposon elements recently amplified and diverged across the eight *Oryza* AA- genomes.

**Figure 3.**
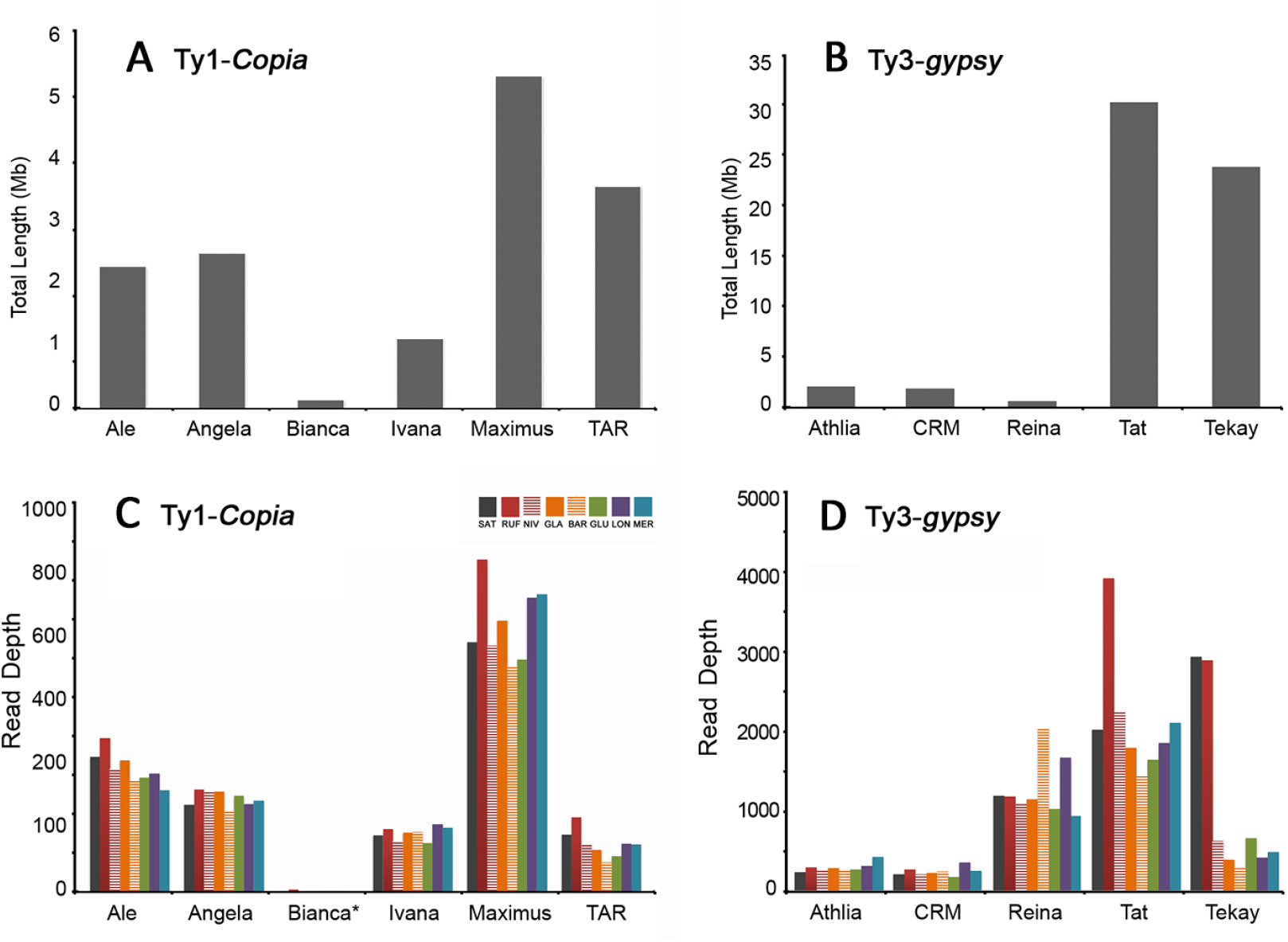
Sequence features of Ty1-*copia* and Ty3-*gypsy* retrotransposon families across the eight AA- genome *Oryza* species. Total sequence of Ty1*-copia* **(A)** and Ty3-*gypsy* **(B)** elements in the SAT genome, annotated using RepeatMasker. LTR reads depth proxy for coy number for various Ty1-*copia* **(C)** and Ty3-*gypsy* **(D)** families.

Phylogenetic analysis reveals that Ty1-*copia* families are more evolutionarily dispersed and smaller in size than Ty3-*gypsy* consistent with previous reports (Vitte *et al.*, 2007). Note that long branches represent early retrotransposon insertions, whereas short clusters indicate new bursts. It is clear that *TAR* possesses a large number of newly generated SAT retrotansposon families, for examples, OAL007 (*Houba*) and OAL013 (*osr1*, *osr5)* (**Figure 2A**). Although the copy number *TAR* was moderate (based on read-depth), the insert length was relatively large in comparisons to other Ty1-*copia* lineages consistent with an increased number of recently amplified intact retroelements (**Figure S2A**; **Figure 3A**). Compared to non-SAT genomes, *Angela* and *Ivana* families drive the latest burst of retrotransposons in SAT. It is interesting to observe that the majority of LTR retrotansposons in *Ale* may represent ancient retrotransposon amplification events as they formed the largest number of long branches in the Ty1-*copia* phylogenetic tree. Both the total length and read depth of the *Maximus* lineage are relatively large, especially in RUF, indicating a substantial contribution to the increase in genome size. In contrast to *Maximus, Bianca* was reported to have become extinct in soybean (Du *et al.*, 2010). This was exemplified by the longest phylogenetic branch lengths (**Figure S2A**), and the shortest insert lengths in the eight rice genomes (**Figure S2A; Figure 3A and B**).

Compared to Ty1-*copia* (Gao *et al.*, 2004), Ty3-*gypsy* retrotransposon elements serve as an important driver of rice genome evolution due to their longer sequence lengths and more recent rounds of amplification. Thus, even though there are fewer Ty3-*gypsy* families than Ty1-*copia*, Ty3-*gypsy* are more prevalent than Ty1-*copia* elements in this set of rice genomes, when compared to SAT alone (McCarthy *et al.*, 2002)and the FF- genome species, *O. brachyantha* (Chen *et al.*, 2013). Phylogenetic analysis not only confirms recent bursts of Ty3-*gypsy* retrotransposon elements in SAT (McCarthy *et al.*, 2002) but also reveals recent amplification of diverse families, usually shown by grouping numerous short branches together across these rice genomes (**Figure 2B**). *Tat* represents such an example and comprises several newly amplified retrotransposon families, such as OAL002, OAL003, OAL005 and OAL008 (**Figure 2B**). The total lengths of *Tat* retroelements are apparently higher than any other Ty3-*gypsy* lineages, probably resulting from high copy number of intact elements (**Figure S2B**; **Figure 3B and D**). *Tekay* typifies the most prevalent group of retrotransposons (e.g., OAL001) (**Figure 2B**, **Figure S2B**; **Figure 3B and D**). New bursts of OAL001 specifically in SAT and RUF are far more abundant than any of the other six rice species (**Figure 2B**; **Figure S2B**; **Fig 3B and D**). *Reina* shows the greatest number of long branches, represented by all eight species, indicating their early integration into the common ancestral genome (**Figure 2B**). Besides, the observation of high LTR retrotransposon copy number (**Figure 3D**) but short insert lengths for *Reina* (**Figure S2B; Figure 3D**) suggests that highly fragmented single-copy elements persist in these rice genomes. The remaining two lineages, *Athila* and *CRM*, show low levels of retrotransposition with both small inset lengths and low numbers of LTR retrotransposons.

Our results show that, in contrast to Ty1-*copia* elements, species-specific bursts of the five Ty3-*gypsy* lineages more frequently occurred and thus more actively driving the genome evolution after recent speciation of these rice species. Rapid amplification of *Tekay* is restricted to SAT and RUF; *Tat* quickly amplified in RUF but was inactive in BAR; recent *Reina* bursts are observed in BAR and LON. As for Ty1-*copia* lineages, only *Maximus* shows evidence of bursts in RUF, LON and MER. By following recent speciation, independent rapid amplifications of LTR retrotransposon lineages have occurred leading to remarkably differing sequence content in these rice genomes. Bursts of *Tat, Tekay* and *Maximum* retrotransposons, for instance, have resulted in an estimated increase of genome size of ~100 Mb in RUF ~0.72 Mya (Zhu *et al.*, 2014), while lineage-specific accumulation of retrotransposons (e.g. *Reina)* has occurred between GLA and its wild progenitor BAR split ~0.26 Mya (Zhang *et al.*, 2014). Moreover, recent bursts of one or more retrotransposon lineages appear to have frequently occurred in specific species: *Tekay* in SAT, *Tat, Tekay* and *Maximum* in RUF, *Reina* in BAR, *Reina* and *Maximum* in LON, and *Maximum* in MER.

### Demographic history of rice retrotransposon families

Comparative analysis of the eight complete rice genomes allow us, for the first time, to trace the life history of retrotransposon families in closely related plant species. Although it is difficult to accurately date the earliest insertion events of a retrotransposon family, the burst periods for each family may be followed by examining the distribution of insertion times (**Figure 1**). Of the top 23 most abundant retrotransposon families in this study, we found that 11 are still active with at least one element having two identical LTRs; the other 12 have completed their entire life histories during an earlier period when AA- genomes diverged. There are typically lower proportions of these elements in SAT when compared to one or more of the other AA- genome species.

From an evolutionary perspective, the accumulation of these retrotransposon families varied dramatically among the lineages (**Figure 1**). Highly amplified retrotransposon families (e.g., OAL005, OAL006, OAL008, OAL010, and OAL012) shared a relatively short half-life when compared to those with fewer retrotransposition events that evolved during similar periods (e.g., OAL015, OAL016, OAL017, OAL018 and OAL019). OAL021 experienced a rapid proliferation of copy number but the shortest life history; nearly all insertions occurred within ~0.2 Myr, approximately equal to when GLU split from the common ancestor of SAT/RUF/NIV and GLA/BAR. The retrotransposition activity of this family declined rapidly during the next 0.2 Myr within the SAT lineage. These results suggest that high levels of retrotransposition activity may be associated with strong negative selection, special environmental stresses or other random events (Grandbastien, 1998; Grandbastien *et al.*, 2005).

In order to understand the evolutionary dynamics of rice LTR retrotransposons in the context of their insertion times, we classified a total of 2,326 intact retroelements from 261 SAT families into high-copy (>20), low-copy (2-20) and single-copy families. Estimation of insertion times suggests that single-copy retrotransposon elements, followed by low-copy families, populated their host genome quite early (~1-10 Mya). The majority of these are incomplete with respect to their LTR retrotransposon structure (**Figure S3A**) but homology searches gleaned a number of retrotransposon fragments in SAT and other seven rice genomes as well. Our calculation of proportions of LTR retrotransposon sequence lengths revealed that high-copy number families possess approximately 2/3 of the total sequence length — far more than low-copy and single-copy number families (**Figure S3B**).

The OAL008 family typifies the evolutionary history of a common LTR retrotransposon across the eight rice species (**Figure S4A**). The normal distribution of insertion times of OAL008 retrotransposons shows no evidence of any new insertions within the last 0.5 Myr; OAL008 came into the host genome and began to amplify before the AA- genome *Oryza* species diverged about ~4.8 Mya. It reached its zenith approximately 1-2 Mya in SAT. The time span from initial insertion to the burst was relatively longer than the period from the burst to the inactivity of retrotransposition. Our data confirm that the half-life of this family is ~ 4 Myr in AA- genome *Oryza* species, which is quite consistent with previous estimates of ~3 to 4 Myr in SAT (Ma and Bennetzen, 2004; Vitte *et al.*, 2007). Phylogenetic analysis of the OAL008 retroelements based on RT alignment of 155 amino acid sequences shows an almost uniform growth of species-specific retrotransposons among these species (**Figure S4A**); LTR retrotransposon copy number analysis indicates that OAL008 was more abundant in RUF and MER than in the other six species (**Figure 1**).

Since the average retrotransposon half-life is ~4 Myr in rice, LTR retrotransposon insertions older than that frequently become highly fragmented, consistent with a pattern of speedy accumulation, decline and extinction. Our study has revealed novel insights into the evolutionary dynamics of retrotransposons: After new retrotransposon lineages are generated and being to integrate into their host genome, some may immediately adopt a normal life-history involving several rounds of burst, accumulation and decline, producing a large number of elements. Others survive and amplify at different rates and then gradually degenerate, or become dormant amplifying at a later date before becoming eliminated. Retrotransposon maintenance and potential is thought to be largely determined by mechanisms such as deletion, unequal recombination, and methylation (Bennetzen, 2002; Petrov *et al.*, 2000; SanMiguel *et al.*, 1998). LTR retrotransposons experience high levels of mutation, rearrangement, and recombination providing a rich genetic resource for the generation of new LTR retrotransposon elements (Dolgin and Charlesworth, 2008; Ma and Bennetzen, 2006). Under conditions of environmental change or especially biotic and abiotic stresses that serve as strong forces of natural selection, some LTR retrotransposons that manage to escape suppression from the host genome may become a new burst branch (Baucom *et al.*, 2009b; Grandbastien, 1998). However, more examples as well as experimental evidence are required to reveal the precise conditions that may stimulate rapid amplifications of some retrotransposon families while suppressing others to produce such a large number of low-copy or single-copy families in host genomes.

### Lineage-specific massive LTR retrotransposon bursts in very recently diverged AA- genome *Oryza* species

The significant correlation of the top 44 most abundant retrotransposon families between SAT and each of the other seven species may indicate that these genomes have experienced a rapid amplification of genome-wide LTR retrotransposons. However, LTR copy number analysis indicates that certain families massively amplified in a lineage-specific manner (**Figure S1B**), and, thus, underwent distinct evolutionary paths since recent split of AA- genome *Oryza* species over the past ~4.8 Myr (Ma and Bennetzen, 2006). To exemplify the idiosyncratic nature of these expansions, we present the findings of the top three most abundant retrotransposon families: OAL001, OAL002 and OAL003.

OAL001 represents the largest family, including the three previously reported retrotransposon families in rice: *RIRE3, RIRE8*, and *Osr34*. A total of 385 OAL001 retrotransposons group into three clusters based on a phylogenetic analysis of LTRs (**Figure 4A**), which is further supported by analysis of the RT sequences (I, II and III) (**Figure 4B**). Detailed analysis of LTR phylogenetic tree shows that this family contains the three Ty3-*gypsy* branches (I, II and III) and one nonautonomous branch (IV). It is clear that Branch IV is derived from Branch III; the nonautonomous branch IV shows the fewest copy number in SAT and possesses highly homologous but longer LTRs from autonomous branches due to insertions (**Figure 4C**). LTR read-depth analysis suggests vast bursts of all four branches of the OAL001 family but only in SAT and RUF and not the other six species (**Figure 4D**). Note that both RUF and NIV are the presumed wild progenitor of SAT; although extensive population sampling of RUF, NIV and SAT is required to further refine the evolutionary dynamics and mechanisms behind this species continuum. Our data support a very recent and massive burst of this largest retrotranposon family immediately after the fairly recent speciation of SAT, NIV and RUF.

**Figure 4.**
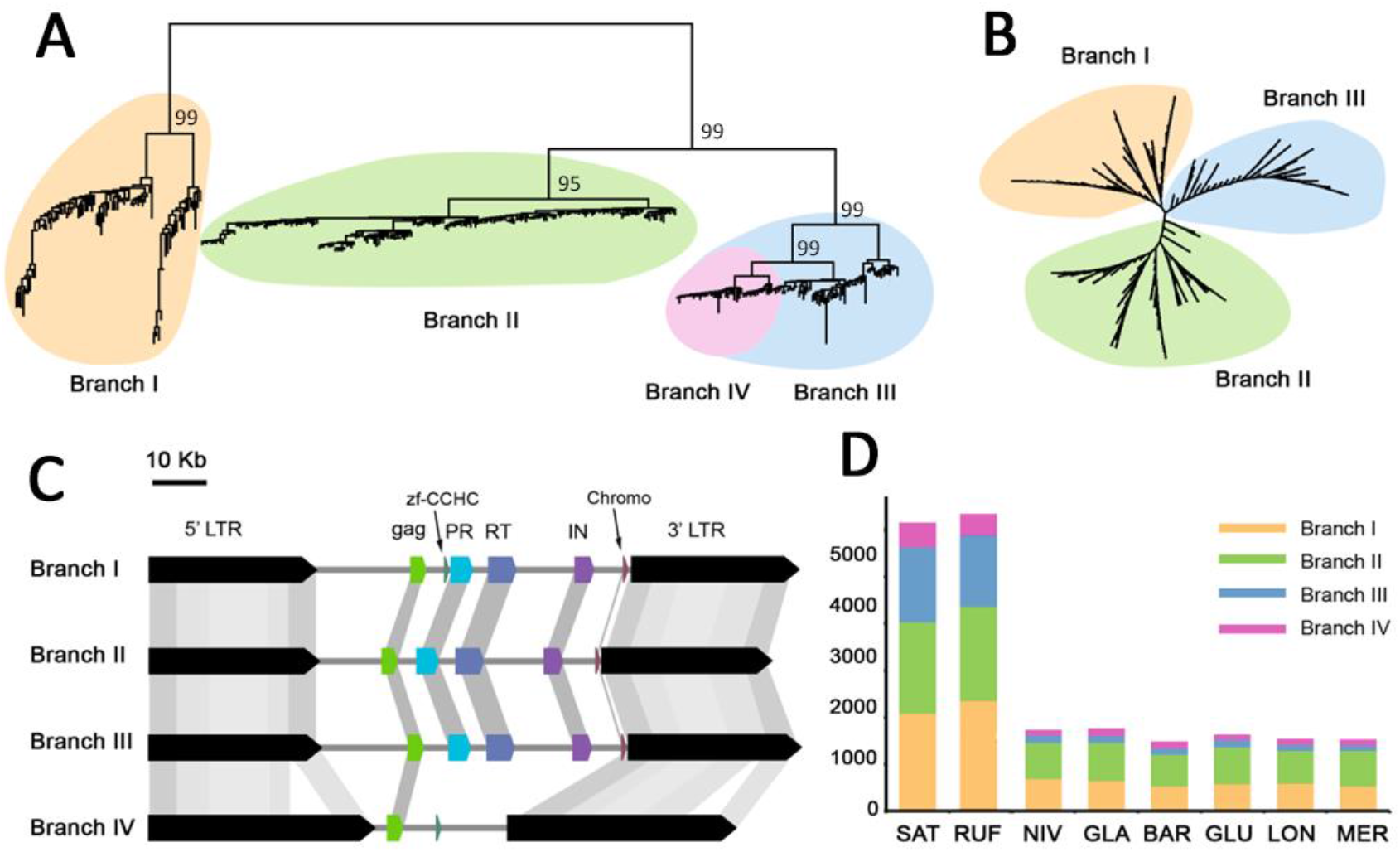
Evolutionary dynamics of the OAL001 family across the eight AA-genome *Oryza* species. **(A)** Phylogram based on LTRs and **(B)** RT sequences; **(C)** genomic structure of retroelements across the 4 branches in OAL001; **(D)** Read depth analysis of each of the four branches in the eight species: Branch I (yellow), Branch II (green), Branch III (blue) and the nonautonomous Branch IV (red).

OAL002 is a Ty3-*gypsy* family formerly known as *hopi* with full length insertions of up to 12 Kb (Picault *et al.*, 2009). Given the relatively long sequence length for each intact element, the growth and decay of a retrotransposon family like OAL002, at least to some extent, has influenced the genome size of AA- *Oryza* species. Phylogenetic analyses of both RT (N= 375) and LTR (N = 373) sequences clearly cluster the retrotransposon elements into two groups (I and II) (**Figure 5A**). Interestingly, the estimation of insertion times suggests that Group I elements are ancient (older than 7 Myr) but experienced only small amplification events after they separated from the other *Tat* families approximately 2.5 to 1 Mya (**Figure 5B**). After a short epoch of silence, massive bursts of retrotransposons (Group II) rapidly occurred in SAT, RUF and NIV around 1 Mya—a time equivalent to the divergence of Asian SAT, RUF and NIV from other AA-genome *Oryza* species (**Figure 5B**). Such an on-going amplification of these three Asian rice species, RUF, SAT and NIV, has contributed large proportion of OAL002 retrotransposons when compared to the other five rice species.

**Figure 5.**
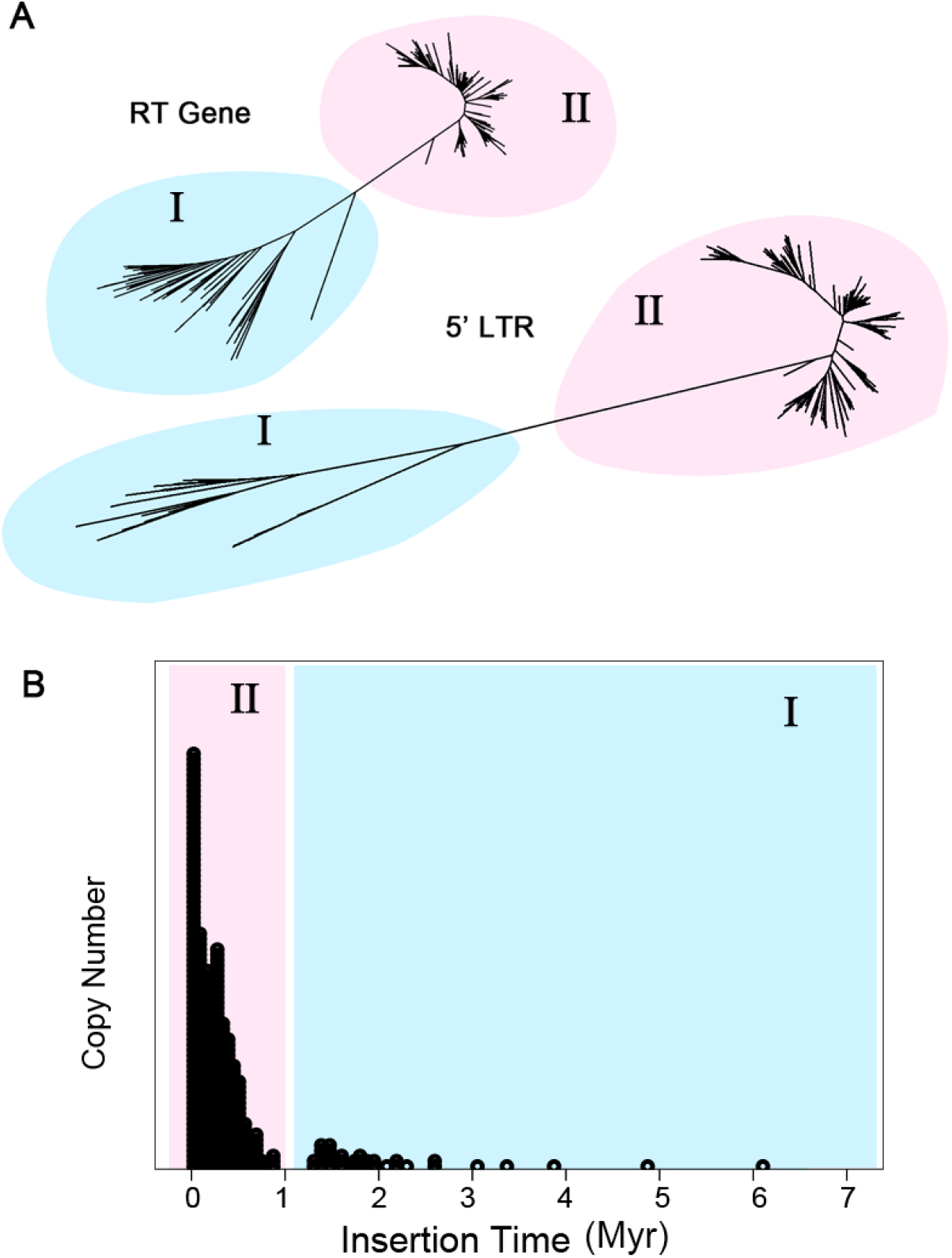
Evolutionary dynamics of the OAL002 family across the eight AA-genome *Oryza* species. **(A)** Phylogentic trees constructed using RT gene and 5’ LTR sequences, respectively. **(B)** Distribution of insertion times. The burst of LTR retrotransposon elements occurred in the Asian lineage (SAT, RUF and NIV) are highlighted (pink) in contrast to those shared by all eight AA- genome *Oryza* species (blue). The burst of LTR retrotransposons is largely restricted to SAT.

OAL003 contains two renowned families, *Dasheng* and *RIRE2*, which have been studied extensively, serving as an excellent model to explore evolutionary relationships between autonomous and nonautonomous retrotransposon elements in plants (Grandbastien *et al.*, 2005; Jiang *et al.*, 2002). Phylogenetic analysis of 930 LTR sequences cluster OAL003 retrotransposons into the eight branches (I, II, III, IV, V, VI, VII and VIII) that separated earlier than the divergence of the eight AA-genome *Oryza* species. Among these, the two most prevalent clades, I and VIII, are equal to *Dasheng* and *RIRE2*, respectively (**Figure 6A**). Previous studies in rice incorporated III, IV, V and VI into an “intermediate” group between *Dasheng* and *RIRE2* using the long branch II as outgroup; these studies also reported that *Dasheng* and *RIRE2* shared similar insertion sites and observed some chimeric *Dasheng/RIRE2* elements (Jiang *et al.*, 2002). In this study, we estimated the number of insertion events and the evolutionary origin of these two groups. Our analysis reveals that the number of nonautonomous *Dasheng* elements has gradually exceeded that of donor *RIRE2* elements over the last 0.5 Myr (**Figure 6B**). This tendency may have limited retrotransposon efficiency by reducing the supply of enzymes needed for a successful retrotransposition. Our results show, in comparison to other AA- genome species, OAL003 retrotransposons became exceptionally amplified in RUF (**Figure 6A and C**). It is apparent that the RUF genome possesses a large quantity of *RIRE2* relative to *Dasheng*, promoting higher reverse transcriptase activity (**Figure 6A**). The mechanisms involved in the enzyme capture and subsequent reverse transcription between *Dasheng* and *RIRE2* still remains unknown. However, the competition between nonautonomous elements and their donors may conceivably explain these potential differences of reverse transcription activity in rice.

**Figure 6.**
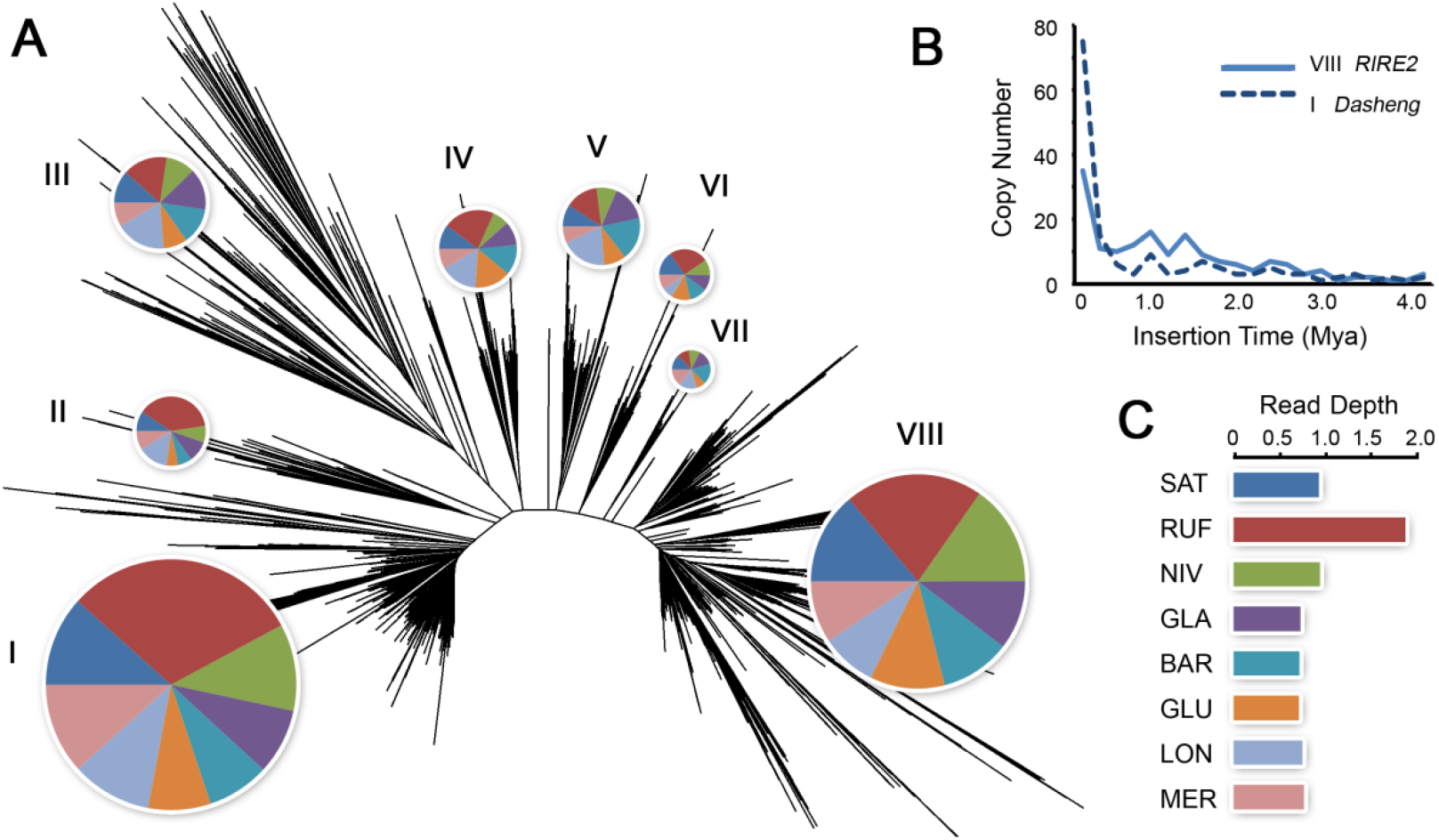
Evolutionary dynamics of the OAL003 family across the eight AA-genome *Oryza* species. **(A)** Radial phylogram clusters OAL003 LTR retrotransposon family into seven clades with the relative proportion by species (see **C** for color key) indicated by the pie chart. The two most prevalent two clades are I (*Dasheng*) and VII (*RIRE2*). **(B)** The insertion times and copy number of clades I (*Dasheng*) and VII (*RIRE2*) in SAT are compared. **(C)** Proportional LTR read depth among the eight species is shown by the colored bars.

On the whole, LTR retrotransposons are the most plentiful in RUF, resulting in the largest genome size among all AA- genome *Oryza* species (**Figure 1B**; **Figure S1**; **Table S2**). Different bursts of retrotransposons also contribute to the slightly enlarged genome sizes of SAT and NIV, which grow almost exclusively in Asia. Although not restricted to SAT, this species has accumulated a number of retrotransposons as a result of especially recent amplifications. Besides the above-described patterns observed in OAL001, OAL002 and OAL003, some species-specific bursts were also observed in SAT (OAL007 and OAL011) and MER (OAL012, OAL014, and OAL023). Almost half of the top 44 most abundant retrotransposon families show high proportions of retrotransposon elements in RUF, followed by SAT, NIV, MER, and LON. In spite of their close relationships, we also observe species-specific retrotransposon differences between GLA and its immediate wild progenitor BAR that diverged merely ~0.26 Mya in Africa (Zhang *et al.*, 2014). It is possible that environmental changes or stochastic mutational processes have induced the species-specific bursts of retrotransposons that previously existed (Grandbastien, 1998; Grandbastien *et al.*, 2005). Our findings are similar to *O. australiensis*, where amplification of only a few LTR retrotransposon families have been sufficient to double its genome size within just a few million years (Piegu *et al.*, 2006).

## Conclusion

The evolutionary dynamics and mechanisms of LTR retrotransposon expansion during speciation are largely unknown. Here, we performed a genome-wide comparative analysis of eight AA- genome *Oryza* species, characterizing a total of 790 LTR retrotransposon families. The resulting evolutionary framework shows that LTR retrotransposons have experienced massive amplifications albeit with fairly divergent and idiosyncratic life histories since these species diverged ~4.8 Myr. This study provides novel insights into the rapid evolution of rice LTR retrotransposons that shaped the architecture and size of rice genomes during and after their recent speciation.

## Acknowledgements

We acknowledge helpful discussions and advice with data sources and software from T. Zhu, Y. Liu, Y. L. Liu, Y. Zhang, E. H. Xia and C. Shi. We are grateful for J.B. Bennetzen for his valuable comments on data analyses. This work was supported by Project of Innovation Team of Yunnan Province, the Top Talents Program of Yunnan Province (20080A009), Hundred Oversea Talents Program of Yunnan Province, Key Project of Natural Science Foundation of Yunnan Province (2010CC011), and Hundred Talents Program of Chinese Academy of Sciences (CAS) (to L. Z. Gao).

## References

Aggarwal RK, Brar DS, Khush GS. 1997. Two new genomes in the Oryza complex identified on the basis of molecular divergence analysis using total genomic DNA hybridization. Molecular and General Genetics MGG 254: 1–12.

Baucom RS, Estill JC, Chaparro C, Upshaw N, Jogi A, Deragon JM, Westerman RP, Sanmiguel PJ, Bennetzen JL. 2009a. Exceptional diversity, non-random distribution, and rapid evolution of retroelements in the B73 maize genome. PLoS Genet 5: e1000732.

Baucom RS, Estill JC, Leebens-Mack J, Bennetzen JL. 2009b. Natural selection on gene function drives the evolution of LTR retrotransposon families in the rice genome. Genome Res 19: 243–254.

Bennetzen JL. 2000. Transposable element contributions to plant gene and genome evolution. Plant Mol Biol 42: 251–269.

Bennetzen JL. 2002. Mechanisms and rates of genome expansion and contraction in flowering plants. Genetica 115: 29–36.

Chen J, Huang Q, Gao D, Wang J, Lang Y, Liu T, Li B, Bai Z, Luis Goicoechea J, Liang C, Chen C, Zhang W, Sun S, Liao Y, Zhang X, Yang L, Song C, Wang M, Shi J, Liu G, Liu J, Zhou H, Zhou W, Yu Q, An N, Chen Y, Cai Q, Wang B, Liu B, Min J, Huang Y, Wu H, Li Z, Zhang Y, Yin Y, Song W, Jiang J, Jackson SA, Wing RA, Wang J, Chen M. 2013. Whole-genome sequencing of *Oryza brachyantha* reveals mechanisms underlying Oryza genome evolution. Nat Commun 4: 1595.

Coffin JM, S. H. Hughes, et al. 1997. Retroviruses.

Cossu RM, Buti M, Giordani T, Natali L, Cavallini A. 2012. A computational study of the dynamics of LTR retrotransposons in the Populus trichocarpa genome. Tree Genetics & Genomes 8: 61–75.

Dolgin ES, Charlesworth B. 2008. The effects of recombination rate on the distribution and abundance of transposable elements. Genetics 178: 2169–2177.

Du J, Tian Z, Hans CS, Laten HM, Cannon SB, Jackson SA, Shoemaker RC, Ma J. 2010. Evolutionary conservation, diversity and specificity of LTR-retrotransposons in flowering plants: insights from genome-wide analysis and multi-specific comparison. Plant J 63: 584–598.

Eickbush TH, Jamburuthugoda VK. 2008. The diversity of retrotransposons and the properties of their reverse transcriptases. Virus Res 134: 221–234.

Estep MC, DeBarry JD, Bennetzen JL. 2013. The dynamics of LTR retrotransposon accumulation across 25 million years of panicoid grass evolution. Heredity (Edinb) 110: 194–204.

Finn RD, Tate J, Mistry J, Coggill PC, Sammut SJ, Hotz HR, Ceric G, Forslund K, Eddy SR, Sonnhammer EL, Bateman A. 2008. The Pfam protein families database. Nucleic Acids Res 36: D281–288.

Fu L, Niu B, Zhu Z, Wu S, Li W. 2012. CD-HIT: accelerated for clustering the next-generation sequencing data. Bioinformatics 28: 3150–3152.

Gao L, McCarthy EM, Ganko EW, McDonald JF. 2004. Evolutionary history of Oryza sativa LTR retrotransposons: a preliminary survey of the rice genome sequences. BMC Genomics 5: 18.

Ge S, Sang T, Lu BR, Hong DY. 1999. Phylogeny of rice genomes with emphasis on origins of allotetraploid species. Proc Natl Acad Sci U S A 96: 14400–14405.

Grandbastien M-A. 1998. Activation of plant retrotransposons under stress conditions. Trends in plant science 3: 181–187.

Grandbastien MA, Audeon C, Bonnivard E, Casacuberta JM, Chalhoub B, Costa AP, Le QH, Melayah D, Petit M, Poncet C, Tam SM, Van Sluys MA, Mhiri C. 2005. Stress activation and genomic impact of Tnt1 retrotransposons in Solanaceae. Cytogenet Genome Res 110: 229–241.

Hřbová E, Neumann P, Matsumoto T, Roux N, Macas J, Doležel J.. 2010. Repetitive part of the banana (*Musa acuminata*) genome investigated by low-depth 454 sequencing. BMC plant biology 10: 204.

Jiang N, Jordan IK, Wessler SR. 2002. Dasheng and RIRE2. A nonautonomous long terminal repeat element and its putative autonomous partner in the rice genome. Plant Physiol 130: 1697–1705.

Jiang SY, Ramachandran S. 2013. Genome-wide survey and comparative analysis of LTR retrotransposons and their captured genes in rice and sorghum. PLoS One 8: e71118.

Jurka J. 1998. Repeats in genomic DNA: mining and meaning. Curr Opin Struct Biol 8: 333–337.

Jurka J. 2000. Repbase update: a database and an electronic journal of repetitive elements. Trends Genet 16: 418–420.

Jurka J, Kapitonov VV, Pavlicek A, Klonowski P, Kohany O, Walichiewicz J. 2005. Repbase Update, a database of eukaryotic repetitive elements. Cytogenet Genome Res 110: 462–467.

Larkin MA, Blackshields G, Brown NP, Chenna R, McGettigan PA, McWilliam H, Valentin F, Wallace IM, Wilm A, Lopez R, Thompson JD, Gibson TJ, Higgins DG. 2007. Clustal W and Clustal X version 2.0. Bioinformatics 23: 2947–2948.

Li R, Yu C, Li Y, Lam TW, Yiu SM, Kristiansen K, Wang J. 2009. SOAP2: an improved ultrafast tool for short read alignment. Bioinformatics 25: 1966–1967.

Li W, Godzik A. 2006. Cd-hit: a fast program for clustering and comparing large sets of protein or nucleotide sequences. Bioinformatics 22: 1658–1659.

Llorens C, Futami R, Covelli L, Dominguez-Escriba L, Viu JM, Tamarit D, Aguilar-Rodriguez J, Vicente-Ripolles M, Fuster G, Bernet GP, Maumus F, Munoz-Pomer A, Sempere JM, Latorre A, Moya A. 2011. The Gypsy Database (GyDB) of mobile genetic elements: release 2.0. Nucleic Acids Res 39: D70–74.

Llorens C, Munoz-Pomer A, Bernad L, Botella H, Moya A.. 2009. Network dynamics of eukaryotic LTR retroelements beyond phylogenetic trees. Biol Direct 4: 41.

Ma J, Bennetzen JL. 2004. Rapid recent growth and divergence of rice nuclear genomes. Proc Natl Acad Sci U S A 101: 12404–12410.

Ma J, Bennetzen JL. 2006. Recombination, rearrangement, reshuffling, and divergence in a centromeric region of rice. Proc Natl Acad Sci U S A 103: 383–388.

Ma J, Devos KM, Bennetzen JL. 2004. Analyses of LTR-retrotransposon structures reveal recent and rapid genomic DNA loss in rice. Genome Res 14: 860–869.

Matyunina LV, Bowen NJ, McDonald JF. 2008. LTR retrotransposons and the evolution of dosage compensation in Drosophila. BMC Mol Biol 9: 55.

McCarthy EM, Liu J, Lizhi G, McDonald JF. 2002. Long terminal repeat retrotransposons of Oryza sativa. Genome Biol 3: 0051–0053.0011.

McCarthy EM, McDonald JF. 2003. LTRSTRUC: a novel search and identification program for LTR retrotransposons. Bioinformatics 19: 362–367.

Meyers BC, Tingey SV, Morgante M. 2001. Abundance, distribution, and transcriptional activity of repetitive elements in the maize genome. Genome Res 11: 1660–1676.

Ouyang S, Buell CR. 2004. The TIGR Plant Repeat Databases: a collective resource for the identification of repetitive sequences in plants. Nucleic Acids Res 32: D360–363.

Pereira V 2004. Insertion bias and purifying selection of retrotransposons in the *Arabidopsis thaliana* genome. Genome Biol 5: R79.

Petrov DA, Sangster TA, Johnston JS, Hartl DL, Shaw KL. 2000. Evidence for DNA loss as a determinant of genome size. Science 287: 1060–1062.

Picault N, Chaparro C, Piegu B, Stenger W, Formey D, Llauro C, Descombin J, Sabot F, Lasserre E, Meynard D. 2009. Identification of an active LTR retrotransposon in rice. The Plant Journal 58: 754–765.

Piednoel M, Carrete-Vega G, Renner SS. 2013. Characterization of the LTR retrotransposon repertoire of a plant clade of six diploid and one tetraploid species. Plant J 75: 699–709.

Piegu B, Guyot R, Picault N, Roulin A, Sanyal A, Kim H, Collura K, Brar DS, Jackson S, Wing RA, Panaud O. 2006. Doubling genome size without polyploidization: dynamics of retrotransposition-driven genomic expansions in Oryza australiensis, a wild relative of rice. Genome Res 16: 1262–1269.

Roulin A, Piegu B, Fortune PM, Sabot F, D’Hont A, Manicacci D, Panaud O 2009. Whole genome surveys of rice, maize and sorghum reveal multiple horizontal transfers of the LTR-retrotransposon Route66 in Poaceae. BMC Evol Biol 9: 58.

SanMiguel P, Gaut BS, Tikhonov A, Nakajima Y, Bennetzen JL. 1998. The paleontology of intergene retrotransposons of maize. Nat Genet 20: 43–45.

Seberg O, Petersen G.. 2009. A unified classification system for eukaryotic transposable elements should reflect their phylogeny. Nat Rev Genet 10: 276.

Smit A, Hubley R, Green P. RepeatMasker Open-3.0. 1996-2010. http://www.repeatmasker.org.

Tamura K, Dudley J, Nei M, Kumar S. 2007. MEGA4: Molecular Evolutionary Genetics Analysis (MEGA) software version 4.0. Mol Biol Evol 24: 1596–1599.

Uozu S, Ikehashi H, Ohmido N, Ohtsubo H, Ohtsubo E, Fukui K. 1997. Repetitive sequences: cause for variation in genome size and chromosome morphology in the genus Oryza. Plant Mol Biol 35: 791–799.

Vaughan DA. 1989. The genu-Oryza L. Currentstatus of taxonomy. IRRI, Manila.

Vaughan DA, Morishima H, Kadowaki K. 2003. Diversity in the Oryza genus. Current opinion in plant biology 6: 139–146.

Vitte C, Panaud O. 2003. Formation of solo-LTRs through unequal homologous recombination counterbalances amplifications of LTR retrotransposons in rice Oryza sativa L. Mol Biol Evol 20: 528–540.

Vitte C, Panaud O. 2005. LTR retrotransposons and flowering plant genome size: emergence of the increase/decrease model. Cytogenet Genome Res 110: 91–107.

Vitte C, Panaud O, Quesneville H. 2007. LTR retrotransposons in rice (*Oryza sativa*, L.): recent burst amplifications followed by rapid DNA loss. BMC Genomics 8: 218.

Wang H, Liu JS. 2008. LTR retrotransposon landscape in Medicago truncatula: more rapid removal than in rice. BMC Genomics 9: 382.

Wawrzynski A, Ashfield T, Chen NW, Mammadov J, Nguyen A, Podicheti R, Cannon SB, Thareau V, Ameline-Torregrosa C, Cannon E, Chacko B, Couloux A, Dalwani A, Denny R, Deshpande S, Egan AN, Glover N, Howell S, Ilut D, Lai H, Del Campo SM, Metcalf M, O’Bleness M, Pfeil BE, Ratnaparkhe MB, Samain S, Sanders I, Segurens B, Sevignac M, Sherman-Broyles S, Tucker DM, Yi J, Doyle JJ, Geffroy V, Roe BA, Maroof MA, Young ND, Innes RW. 2008. Replication of nonautonomous retroelements in soybean appears to be both recent and common. Plant Physiol 148: 1760–1771.

Wessler SR, Bureau TE, White SE. 1995. LTR-retrotransposons and MITEs: important players in the evolution of plant genomes. Curr Opin Genet Dev 5: 814–821.

Wicker T, Keller B. 2007. Genome-wide comparative analysis of *copia* retrotransposons in Triticeae, rice, and *Arabidopsis* reveals conserved ancient evolutionary lineages and distinct dynamics of individual copia families. Genome Res 17: 1072–1081.

Xiong Y, Eickbush TH. 1990. Origin and evolution of retroelements based upon their reverse transcriptase sequences. EMBO J 9: 3353–3362.

Yang Z. 2007. PAML 4: phylogenetic analysis by maximum likelihood. Mol Biol Evol 24: 1586–1591.

Zhang QJ, Zhu T, Xia EH, Shi C, Liu YL, Zhang Y, Liu Y, Jiang WK, Zhao YJ, Mao SY, Zhang LP, Huang H, Jiao JY, Xu PZ, Yao QY, Zeng FC, Yang LL, Gao J, Tao DY, Wang YJ, Bennetzen JL, Gao LZ. 2014. Rapid diversification of five Oryza AA genomes associated with rice adaptation. Proc Natl Acad Sci U S A 111: E4954–4962.

Zhang Y, Liu XS, Liu Q-R, Wei L. 2006. Genome-wide in silico identification and analysis of cis natural antisense transcripts (cis-NATs) in ten species. Nucleic acids research 34: 3465–3475.

Zhu Q, Ge S. 2005. Phylogenetic relationships among A-genome species of the genus Oryza revealed by intron sequences of four nuclear genes. New Phytol 167: 249–265.

Zhu T, Xu PZ, Liu JP, Peng S, Mo XC, Gao LZ. 2014. Phylogenetic relationships and genome divergence among the AA- genome species of the genus Oryza as revealed by 53 nuclear genes and 16 intergenic regions. Mol Phylogenet Evol 70: 348–361.

